# Regulation of stress-induced sleep fragmentation by preoptic glutamatergic neurons

**DOI:** 10.1101/2022.11.30.518589

**Authors:** Jennifer Smith, Adam Honig-Frand, Hanna Antila, Kevin Beier, Franz Weber, Shinjae Chung

**Author notes:** Correspondence: Shinjae Chung, Address: 10-133 Smilow Center for Translational Research, 3400 Civic Center Blvd. Bldg. 421 Philadelphia, PA 19104-6168.

## Abstract

Sleep disturbances are detrimental for our behavioral and emotional well-being. Stressful events disrupt sleep, in particular by inducing brief awakenings (microarousals, MAs) resulting in sleep fragmentation. The preoptic area of the hypothalamus (POA) is crucial for sleep control. However, how POA neurons contribute to the regulation of MAs and thereby impact sleep quality is unknown. Using fiber photometry recordings in mice, we examined the activity changes of genetically defined POA subpopulations during sleep. We found that POA glutamatergic neurons are rhythmically activated in synchrony with an infraslow rhythm in the spindle band of the electroencephalogram during non-rapid eye movement sleep (NREMs) and are transiently activated during MAs. Optogenetic stimulation of these neurons strongly promotes MAs. Exposure to acute social defeat stress significantly increased the number of transients in the calcium activity of POA glutamatergic neurons during NREMs. Optogenetic inhibition during spontaneous sleep and after stress reduced MAs during NREMs and consequently consolidated sleep. Monosynaptically-restricted rabies tracing revealed that POA glutamatergic neurons are innervated by brain regions regulating stress and sleep. Our findings uncover a novel circuit mechanism by which POA excitatory neurons regulate sleep quality after stress.

## INTRODUCTION

Stress disturbs sleep. In particular, sleep fragmentation is one of the key symptoms in stress-related sleep disorders such as insomnia and post-traumatic stress disorders (PTSD) (Germain et al., 2008; Germain, 2013; Van Someren, 2020). Fragmented sleep due to an increased number of MAs impairs memory and induces anxiety, demonstrating the importance of consolidated sleep (McCoy et al., 2007; Tartar et al., 2009; Van Der Werf et al., 2009; Rolls et al., 2011). However, the mechanisms underlying the regulation of MAs and how they are affected by stress remains largely unknown. MAs are brief arousal-like bouts (< 20 s) that occur during NREMs and are characterized by a desynchronized electroencephalogram (EEG) together with an activated electromyography (EMG) in rodents (Halász et al., 1979, 2004; Watson et al., 2016; dos Santos Lima et al., 2019; Antila et al., 2022). MAs preferentially occur at the trough of an infraslow rhythm in the EEG σ (10.5-16 Hz) power which has been shown to regulate transitions from NREMs to MAs or to REMs (Lecci et al., 2017; Cardis et al., 2021; Stucynski et al., 2022; Antila et al., 2022; Kjaerby et al., 2022). Stress has been shown to disrupt the infraslow rhythm as well as the coupling of MAs to this rhythm (Antila et al., 2022).

The POA is crucial for sleep regulation (Sherin et al., 1996; Szymusiak et al., 1998; Chung et al., 2017; Kroeger et al., 2018; Ma et al., 2019; Vanini et al., 2020). Stress-induced insomnia in rats led to increased levels of c-Fos in POA neurons (Cano et al., 2008). However, the identity of the activated neurons and their dynamics during sleep are still unknown. Physical stress activates POA glutamatergic neurons (Zhang et al., 2021), however their role in mediating stress-related sleep disturbances is not yet characterized. By employing an acute social defeat stress paradigm together with fiber photometry and optogenetic manipulation, we investigated the role of the POA in MA regulation during spontaneous sleep and following stress.

## RESULTS

### The activity of POA glutamatergic neurons rhythmically fluctuates during NREMs and coincides with MAs

We hypothesized that POA neurons with strong modulation in their activity during MAs are likely to be involved in sleep fragmentation. As MAs coincide with the trough of the infraslow oscillation in the EEG σ power (Lecci et al., 2017; Cardis et al., 2021; Antila et al., 2022; Kjaerby et al., 2022), we predicted that the activity of POA subpopulations that regulate MAs is closely correlated with this infraslow σ rhythm. We set out to systematically examine the activity dynamics of multiple genetically defined POA subpopulations during sleep, including glutamatergic and GABAergic neurons as well as cholecystokinin (CCK) and tachykinin 1 (TAC1) neurons that have been previously shown to be sleep-promoting (Chung et al., 2017). To monitor the dynamics of genetically defined POA subpopulations during spontaneous sleep, we injected Cre-dependent adeno-associated viruses encoding the calcium indicator GCaMP6s (AAVs-Syn-Flex-GCaMP6s) into the POA of VGLUT2-, VGAT-, CCK- and TAC1-Cre mice and implanted an optic fiber to measure the calcium-dependent fluorescence using fiber photometry together with EEG and EMG recordings (**Figures 1A and S1**). During NREMs, we found that POA VGLUT2 neurons become activated during MAs, whereas the activity of VGAT, CCK, and TAC1 neurons was not significantly altered (**Figures 1B, 1D and S2**; one-way repeated measures [rm] ANOVA P = 1.542e-20, 0.361, 0.993, 0.998 for VGLUT2, VGAT, CCK and TAC1). The calcium signal of POA VGLUT2 neurons started increasing 0.5 s before the onset of MAs (**Figure 1D**; one-way rm ANOVA followed by pairwise t-tests with Bonferroni correction, P = 0.043). 32.47 ± 2.25 % (mean ± s.e.m.) of the POA VGLUT2 calcium transients during NREMs overlapped with MAs (**Figure 1B**). During the remaining calcium transients, NREMs was not interrupted. The calcium signal was significantly higher, when it coincided with a MA (**Figure 1E**, paired t test, P = 1.916e-15).

**Figure 1.**
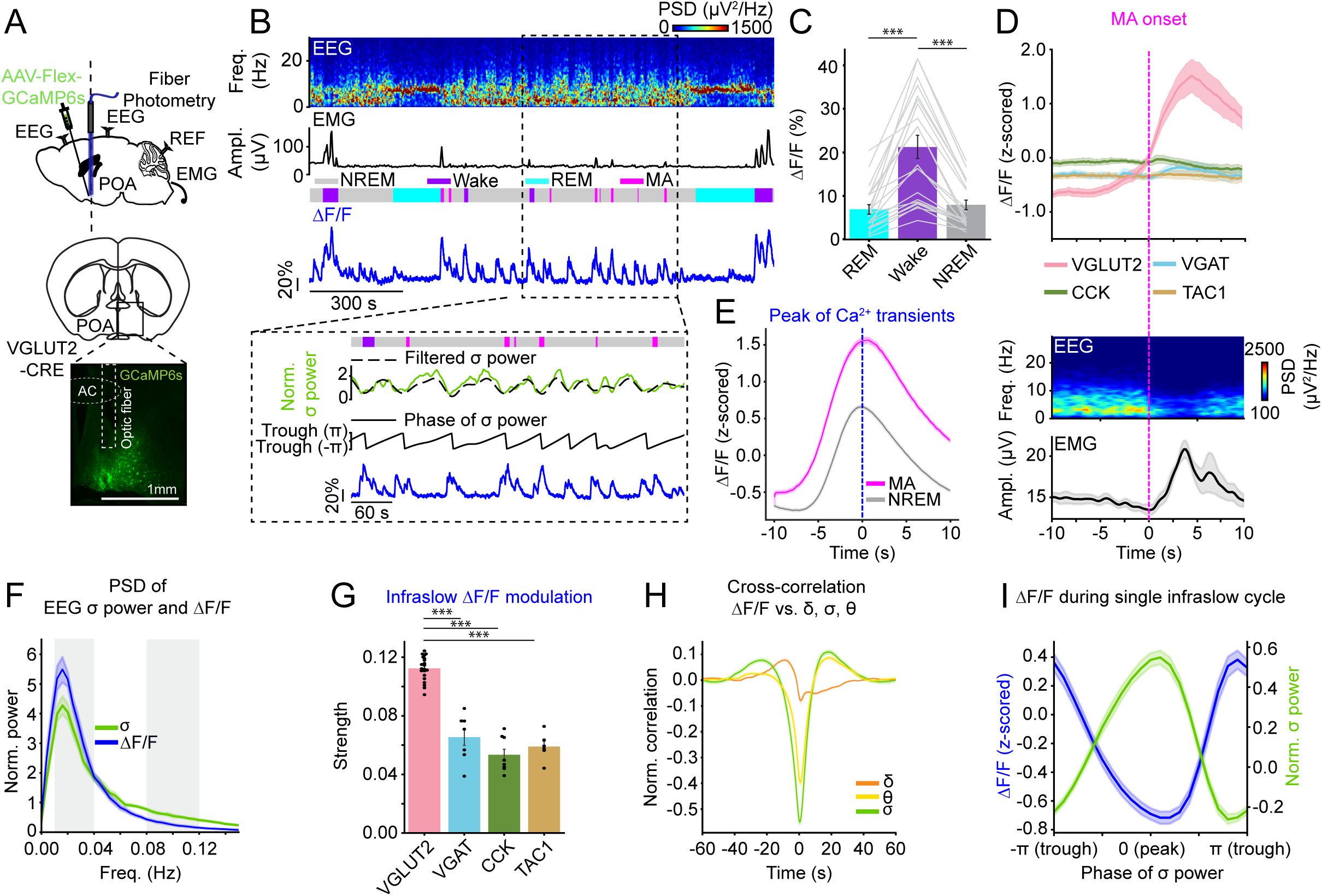
POA VGLUT2 neurons become rhythmically activated during NREMs. **(A)** Schematic of fiber photometry with simultaneous EEG and EMG recordings in different POA subpopulations. Mouse brain figure adapted from Allen mouse brain atlas (© 2015 Allen Institute for Brain Science). Bottom, fluorescence image of POA in a VGLUT2-Cre mouse injected with AAV-FLEX-GCaMP6s into the POA. Scale bar, 1 mm. **(B)** Top, example fiber photometry recording. Shown are parietal EEG spectrogram, EMG amplitude, color-coded brain states, and ΔF/F signal. Bottom, brain states, EEG σ power (10.5 - 16 Hz, green, parietal EEG), phase of EEG σ power, and calcium signal during a selected interval (dashed box) at an expanded timescale. The filtered σ power (dashed line) was used to determine the phase of the σ power oscillation (middle). Freq, frequency. **(C)** ΔF/F activity during REMs, wake, and NREMs. Bars, averages across mice; lines, individual mice; error bars, ± s.e.m. One-way rm ANOVA followed by pairwise t tests with Bonferroni correction, ***P < 0.001. n = 21 mice. **(D)** Calcium activity changes in different POA subpopulations, EEG spectrogram, and EMG amplitude at the transition to MAs. n = 21 (VGLUT2), 7 (VGAT), 8 (CCK) and 6 (TAC1) mice. **(E)** Calcium activity of VGLUT2 neurons during MAs and continuous NREMs episodes. Time point 0 s corresponds to the peak of calcium transients. Paired t test, ***P < 0.001. **(F)** Normalized power spectral density (PSD) of EEG σ power and calcium activity during NREMs. **(G)** Strength of the infraslow calcium activity oscillation. To quantify the strength of the infraslow rhythm, we computed the areas under the PSD in the ranges 0.01 - 0.04 Hz and 0.08 - 0.12 Hz respectively (gray boxes in Figure 1F), and subtracted the second value from the first value. Bars, averages across mice; dots, individual mice; error bars, ± s.e.m. One-way ANOVA followed by t tests with Bonferroni correction, ***P < 0.001. **(H)** Cross-correlation of ΔF/F activity with EEG δ, θ and σ power. **(I)** Average calcium activity during a single cycle of the σ power oscillation. **(D, E, F, H, I)** Shadings, ± s.e.m.

**Figure 2.**
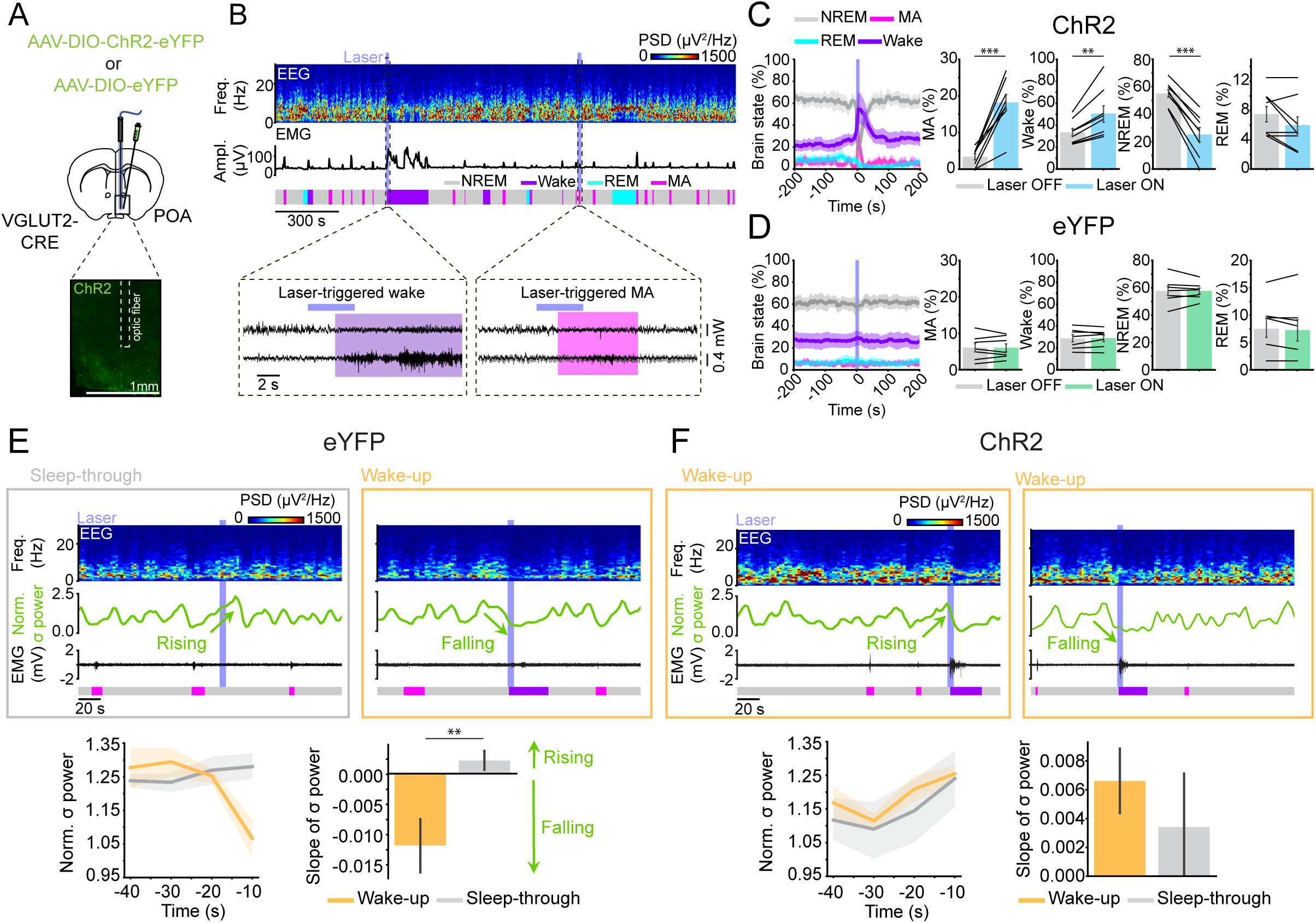
Activation of POA VGLUT2 neurons promotes arousal. **(A)** Schematic of optogenetic activation experiments. Bottom, fluorescence image of POA in VGLUT2-CRE mouse injected with AAV-DIO-ChR2-eYFP into the POA. Scale bar, 1 mm. **(B)** Example laser session. Shown are parietal EEG spectrogram, EMG amplitude, color-coded brain states, and EEG and EMG raw traces during selected periods (dashed boxes) on an expanded timescale. Blue shading, laser stimulation interval (10 Hz, 4 s). **(C)** Percentage of MAs, wake, NREMs, or REMs before and during laser stimulation (10 Hz, 4 s) in ChR2 mice. The duration of the baseline interval without laser was equal to that of the following laser interval. Shadings, 95% confidence intervals (CIs). Error bars, ± s.e.m. Paired t tests, ***P < 0.001; **P < 0.01. n = 8 mice. **(D)** Percentage of MAs, wake, NREMs, or REMs before and during laser stimulation (10 Hz, 4 s) in eYFP control mice. Shadings, 95% CIs. Error bars, ± s.e.m. n = 7 mice **(E)** Top, Example sleep-through trial with rising EEG σ power (left) and wake-up trial with falling EEG σ power (right) in eYFP mouse. Shown are parietal EEG spectrograms, EEG σ power, EMG raw traces and color-coded brain states. Blue shading, laser stimulation interval (10 Hz, 4 s). Bottom, EEG σ power (left) and its slope (right) before wake-up and sleep-through trials. Slope is calculated between the σ powers from −30 sec to 0 sec before the laser onset. Shadings and error bars, ± s.e.m. T test, **P < 0.01. n = 7 mice. **(F)** Example wake-up trials in ChR2 mouse. Bottom, EEG σ power (left) and its slope (right) before wake-up and sleep-through trials. Shadings and error bars, ± s.e.m. n = 8 mice.

During NREMs, the POA VGLUT2 neuron activity showed strong fluctuations on the ~minute time scale (**Figure 1B**). We tested whether these fluctuations are correlated with the infraslow rhythm in the EEG σ power. We found that the peaks of power spectral densities (PSDs) from both the ΔF/F signal and the σ power closely overlapped (**Figure 1F**; 0.017 ± 0.004 and 0.017 ± 0.003, respectively). To quantify how strongly the ΔF/F signal is modulated on the infraslow timescale, we determined the ratio of the area under the PSD curve in the ranges of 0.01 - 0.04 Hz and 0.08 - 0.12 Hz (**Figure 1F**). The larger the ratio, the stronger the calcium signal was dominated by fluctuations in the infraslow range (0.01 - 0.04 Hz). The strength of the infraslow fluctuation in the calcium activity was significantly higher for POA VGLUT2 neurons compared with all other cell types (**Figure 1G**; one-way ANOVA, P = 4.551e-17; t-tests with Bonferroni correction, P = 1.811e-10, 1.276e-14, 5.382e-13 for VGLUT2 vs. VGAT, CCK or TAC1). The ΔF/F signal was negatively correlated with both the σ and θ power (**Figures 1H and 1I**). Compared with the δ and θ power, the correlation was strongest for the σ power (**Figure 1H**; one-way rm ANOVA, P = 7.399e-34, pairwise t-tests with Bonferroni correction, P = 3.145e-20, 4.352e-12 for σ vs. δ and σ vs. θ power).

Another striking difference between the different POA subpopulations was their state-dependent activity. POA VGLUT2 neurons were most active during wakefulness and less active during NREMs and REMs (**Figure 1C**; one-way rm ANOVA, P = 1.665e-7; pairwise t-tests with Bonferroni correction, P = 1.000, 9.655e-7, 1.170e-5, for NREM vs. REM, NREM vs. wake and REM vs. wake). In contrast, VGAT, CCK, and TAC1 neurons were most active during REMs (**Figure S2**; one-way rm ANOVA, P = 7.094e-6, 0.001, 2.422e-4 for VGAT, CCK and TAC1; pairwise t-tests with Bonferroni correction, VGAT P = 0.001, 0.014, 0.005, CCK P = 0.016, 0.115, 0.034, TAC1 P = 0.009, 0.020, 0.036 for NREM vs. REM, NREM vs. wake and REM vs. wake, respectively).

Taken together, the activity of POA glutamatergic neurons rhythmically fluctuated during NREMs in synchrony with the infraslow σ rhythm, and transients in their calcium activity often occurred during MAs providing correlational evidence for a role in MA regulation.

### Activation of POA glutamatergic neurons strongly promotes wakefulness

To test whether the POA VGLUT2 neurons promote arousal, we optogenetically activated these neurons for short periods of time (4 sec). VGLUT2-Cre mice were injected with AAVs encoding Cre-dependent channelrhodopsin 2 (AAV-DIO-ChR2-eYFP) into the POA (**Figure 2A**). Stimulating POA VGLUT2 neurons (10 Hz for 4 sec) significantly increased the percentage of MAs and wake during the laser interval while decreasing NREMs (**Figures 2B and 2C**; Paired t-tests, P = 1.824e-4, 0.004, 7.312e-6 for MA, wake and NREMs). In control mice expressing eYFP in the POA VGLUT2 neurons, laser stimulation did not significantly affect the brain state (**Figure 2D**; Paired t-tests, P = 0.935, 0.748, 0.969, 0.5 for MA, Wake, NREMs and REMs).

MAs preferentially occur during the falling phase of the EEG σ power, when mice are more susceptible to wake up in response to acoustic noise (Lecci et al., 2017; Cardis et al., 2021; Antila et al., 2022; Kjaerby et al., 2022). In contrast, mice are less prone to wake up during the rising phase of the infraslow σ rhythm. To test whether optogenetic stimulation-induced arousals depend on the phase of infraslow σ rhythm, we examined the EEG σ power before the optogenetic onset, and split the trials into wake-up and sleep-through trials, depending on whether the animals woke up or not in response to the optogenetic stimulation. In eYFP mice, prior to wake-up trials, the σ power was significantly declining compared to sleep-through trials (**Figure 2E**; Left, Mixed ANOVA, response P = 0.413, time P = 0.054, interaction P = 0.015; Right, t-test for slope of σ power, P = 0.005). This is similar to the declining σ power and decreased number of sleep spindles before the onset of spontaneous MAs (Lecci et al., 2017; Antila et al., 2022; Kjaerby et al., 2022). In contrast, in ChR2 mice, the σ power before laser was not significantly different during optogenetic stimulation induced wake-up and sleep-through trials (**Figure 2F**; Left, Mixed ANOVA, response P = 0.510, time P = 0.145, interaction P = 0.919; Right, t-test for slope of σ power, P = 0.465), suggesting that stimulating POA VGLUT2 neurons induced arousals regardless of the phase of the infraslow σ rhythm. Taken together, activation of POA VGLUT2 neurons powerfully induce arousal during NREMs regardless of the infraslow σ rhythm.

### POA glutamatergic neurons become frequently activated during NREMs after stress

To investigate whether the activity of POA VGLUT2 neurons contributes to sleep fragmentation after stress, we performed fiber photometry recordings together with an acute social defeat stress paradigm (**Figure 3A**). We first performed fiber photometry baseline recordings of POA VGLUT2 neurons. On the stress day, the experimental mouse was exposed to an aggressive CD1 mouse for physical attacks and subsequently stayed in the cage of the CD1 mouse (with the CD1 mouse removed from the cage). Similar to our previous report, acute social defeat stress significantly increased the time spent in wakefulness while decreasing NREMs and REMs (**Figure S3**; paired t-tests, P = 0.008, 0.020, 1.079e-5 for Wake, NREMs and REMs) (Antila et al., 2022). During NREMs, the number of MAs was increased and the duration of NREMs episodes was decreased, resulting in a fragmentation of NREMs (**Figures 3B and S3B**; paired t-tests, P = 1.345e-4, 6.854e-5 for MAs and NREMs duration). The number of sleep spindles and the strength of the infraslow modulation in the σ power was significantly decreased suggesting that acute social defeat stress weakens the infraslow σ rhythm (**Figures 3C and 3D**; paired t-tests, P = 1.422e-4, 0.015 for spindles and strength). Analyzing the activity of POA VGLUT2 neurons, we found that the number of calcium transients was significantly increased during NREMs, and the strength of the infraslow modulation in the ΔF/F signal was decreased (**Figures 3E and 3F**; paired t-tests, P = 0.011, 0.023 for calcium transients and strength).

**Figure 3.**
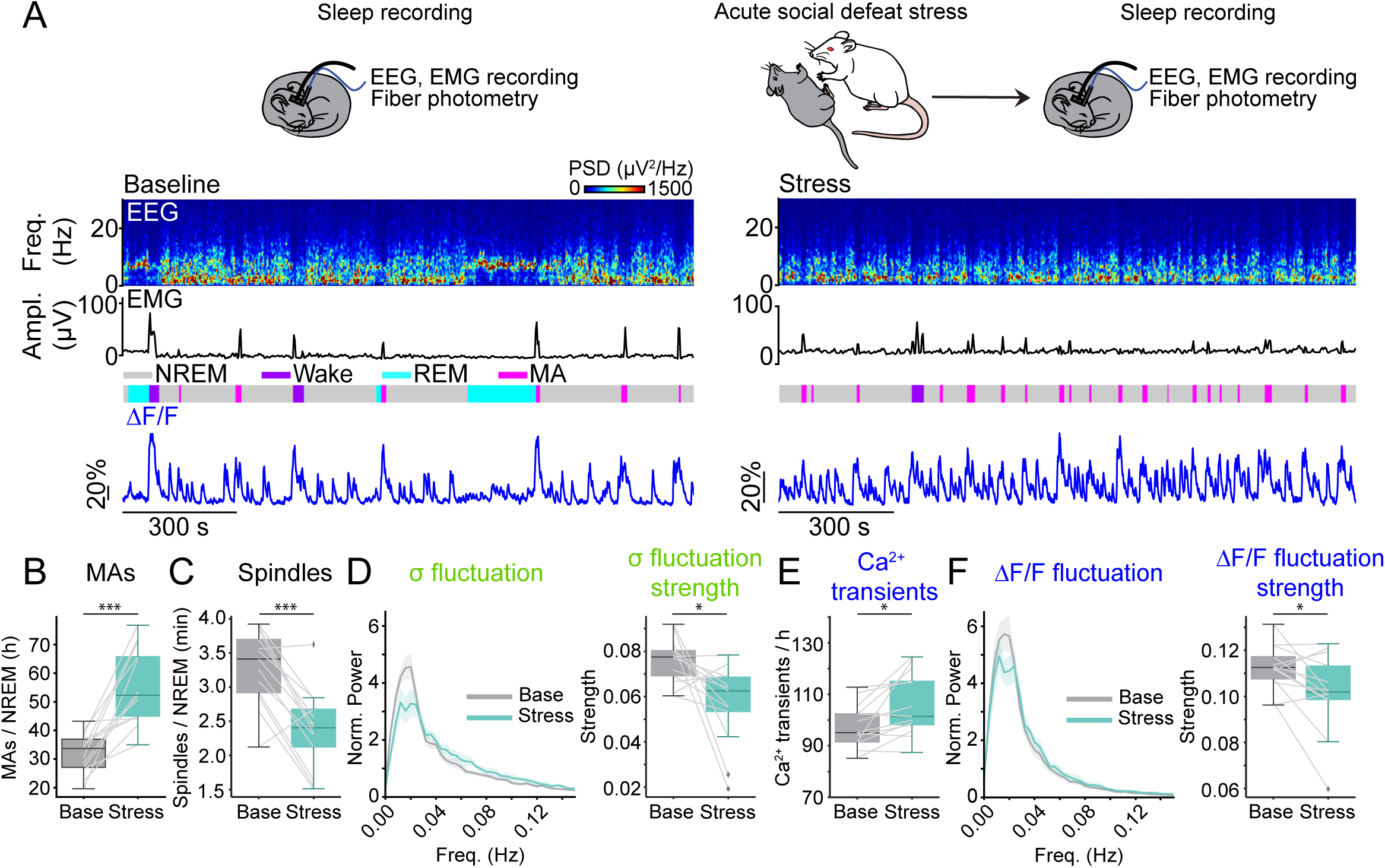
Acute social defeat stress leads to frequent activation of POA VGLUT2 neurons during NREMs. **(A)** Top, Schematic of fiber photometry during baseline recordings and after acute social defeat stress. Bottom, Fiber photometry recordings during baseline recordings (left) and after acute social defeat stress (right). Shown are EEG power spectra, EMG amplitude, color-coded brain states, and ΔF/F signal. **(B)** Number of MAs during NREMs after stress. **(C)** Number of sleep spindles. **(D)** PSD of EEG σ power fluctuations and its strength. **(E)** Number of calcium transients during NREMs. **(F)** PSD of calcium activity fluctuations during NREMs and its strength. **(B-F)** Box plots; gray lines, individual mice. Shadings, 95% CIs. Paired t-tests, ***P < 0.001; *P < 0.05. n = 13 mice.

### Inhibiting POA glutamatergic neurons attenuates stress-induced NREMs fragmentation

To test the necessary role of the activity of POA VGLUT2 neurons in regulating MAs during spontaneous sleep and after stress, we bilaterally injected AAVs encoding the bistable chloride channel SwiChR++ (AAV-EF1α-DIO-SwiChR++-eYFP) (Berndt et al., 2016) into the POA of VGLUT2-Cre mice to optogenetically inactivate POA VGLUT2 neurons for an extended period of time (**Figures 4A-C**). We compared the number of MAs and the percentage, duration, and frequency of each state in 4 hr baseline recordings with sustained laser stimulation (3 s step pulses at 1 m intervals for 4 hrs) between SwiChR++ and control animals expressing eYFP.

**Figure 4.**
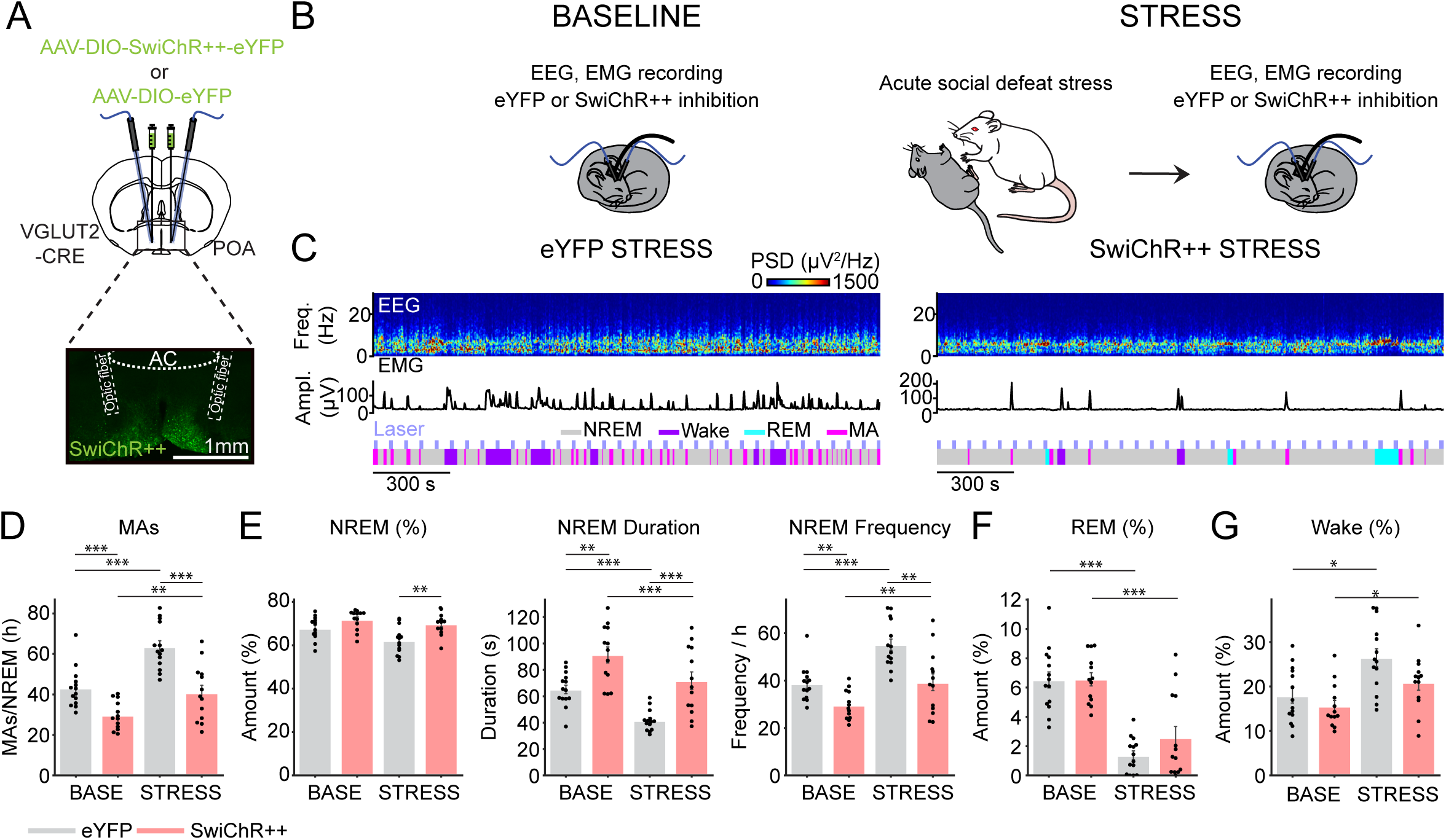
Inhibiting POA VGLUT2 neurons decreased stress-induced fragmentation of NREMs. **(A)** Schematic of SwiChR++-mediated inhibition experiments. Bottom, fluorescence image of POA in a VGLUT2-Cre mouse injected with AAV-DIO-SwiChR++-eYFP bilaterally into the POA. Scale bar, 1 mm. **(B)** Schematic of optogenetic inhibition during baseline recordings and after acute social defeat stress. **(C)** Example session from eYFP (left) or SwiChR++ mouse (right) with laser stimulation after stress. Shown are parietal EEG power spectra, EMG amplitude, and color-coded brain states. **(D)** Number of MAs during NREMs in eYFP (n = 14) and SwiChR++ (n = 13) mice during the 4 hr laser baseline recordings and after stress. **(E)** Percentage of time spent in NREMs, duration and frequency of NREMs episodes during laser baseline and stress recordings. **(F)** Percentage of time spent in REMs. **(G)** Percentage of time spent in wakefulness. **(D-G)** Bars, averages across mice; dots, individual mice; error bars, ± s.e.m. Mixed ANOVA followed by pairwise t-tests with Bonferroni correction, ***P < 0.001; **P < 0.01; *P < 0.05.

Compared with eYFP animals, SwiChR++-mediated inhibition decreased the number of MAs and prolonged NREMs episodes without changing the time spent in NREMs (**Figures 4D and 4E**; Mixed ANOVA, virus P = 3.601e-5, 3.437e-4, 0.0002, stress P = 4.376e-8, 0.013, 1.219e-8, interaction P = 0.033, 0.224, 0.455; t-tests with Bonferroni correction, eYFP-base vs. SwiChR-base, P = 0.0009, 0.085, 0.002 for MAs, NREMs % and NREMs duration respectively). As observed above for the photometry recordings, stress significantly increased the number of MAs and consequently reduced the NREMs episode duration in eYFP mice (**Figures 4D and 4E**; eYFP-base vs. eYFP-stress P = 0.00003, 4.378e-5 for MAs and NREMs duration). SwiChR++-mediated inhibition of POA VGLUT2 neurons significantly lowered the number of MAs compared with that in eYFP mice after stress and consequently increased the duration and decreased the frequency of NREMs episodes (**Figures 4D and 4E**; eYFP-stress vs. SwiChR-stress, P = 1.68e-4, 5.153e-4, 0.003 for MAs, NREMs duration and frequency). The amount of time spent in NREMs was slightly increased by inhibition of POA VGLUT2 neurons after stress (**Figure 4E**, eYFP-stress vs. SwiChR-stress P = 0.003). REMs and wake was not altered by inhibition of POA VGLUT2 neurons during baseline sleep and after stress (**Figures 4F, 4G and S4**; Mixed ANOVA, virus P = 0.027, 0.367, stress P = 5.523e-4, 2.174e-11, interaction P = 0.373, 0.160; t-tests with Bonferroni correction, eYFP-base vs. SwiChR-base P = 0.576, 1.000, eYFP-stress vs. SwiChR-stress P = 0.098, 0.320 for Wake % and REMs %). Taken together, the activity of POA VGLUT2 neurons regulates MAs and thereby influences sleep fragmentation during spontaneous sleep and after stress.

### Presynaptic inputs to POA glutamatergic neurons

The POA is known to receive inputs from multiple brain regions (Chou et al., 2002). To identify presynaptic input neurons that innervate the POA VGLUT2 neurons, we used mono-synaptically restricted rabies tracing (Beier et al., 2015). AAVs with Cre-dependent expression of TVA receptor fused with mCherry and rabies glycoprotein (RG) were injected into the POA (**Figure 5A**). Two weeks later, a modified rabies virus expressing GFP (RV*d*G-GFP) was injected into the POA. The majority of starter cells expressing both TVA-mCherry and GFP was located in the POA. Input neurons were found in the hypothalamus, in particular paraventricular nucleus (PVN), known to be strongly involved in stress responses (Herman and Cullinan, 1997) (**Figure 5B**). Furthermore, POA VGLUT2 neurons receive inputs from the lateral hypothalamus (LH), dorsomedial nucleus of the hypothalamus (DMH), zona incerta (ZI) and arcuate hypothalamic nucleus (ARH), areas known to be involved in the regulation of sleep and wakefulness as well as feeding (Liu et al., 2017; Yamashita and Yamanaka, 2017; Chen et al., 2018; Goldstein et al., 2018; Arrigoni et al., 2019). Monosynaptic input neurons were also found in the lateral septum (LS), medial amygdala (MEA) and piriform cortex (PIR) that are known to regulate stress responses (Nacher et al., 2004; Singewald et al., 2011; Kondoh et al., 2016; Raam and Hong, 2021). Furthermore, POA VGLUT2 neurons are innervated by the subfornical organ (SFO), known to regulate fluid homeostasis, the periaqueductal gray (PAG) in the midbrain involved in the regulation of pain and sleep and the parabrachial nucleus (PBN), involved in thermoregulation (Behbehani, 1995; Weber et al., 2018; Zimmerman et al., 2019; Norris et al., 2021) suggesting that these physiological functions are tightly connected to sleep and wake regulatory neurons in the POA. Therefore, these presynaptic monosynaptic neurons may modulate the POA VGLUT2 neuron activity and regulate the stability of sleep after stress.

**Figure 5.**
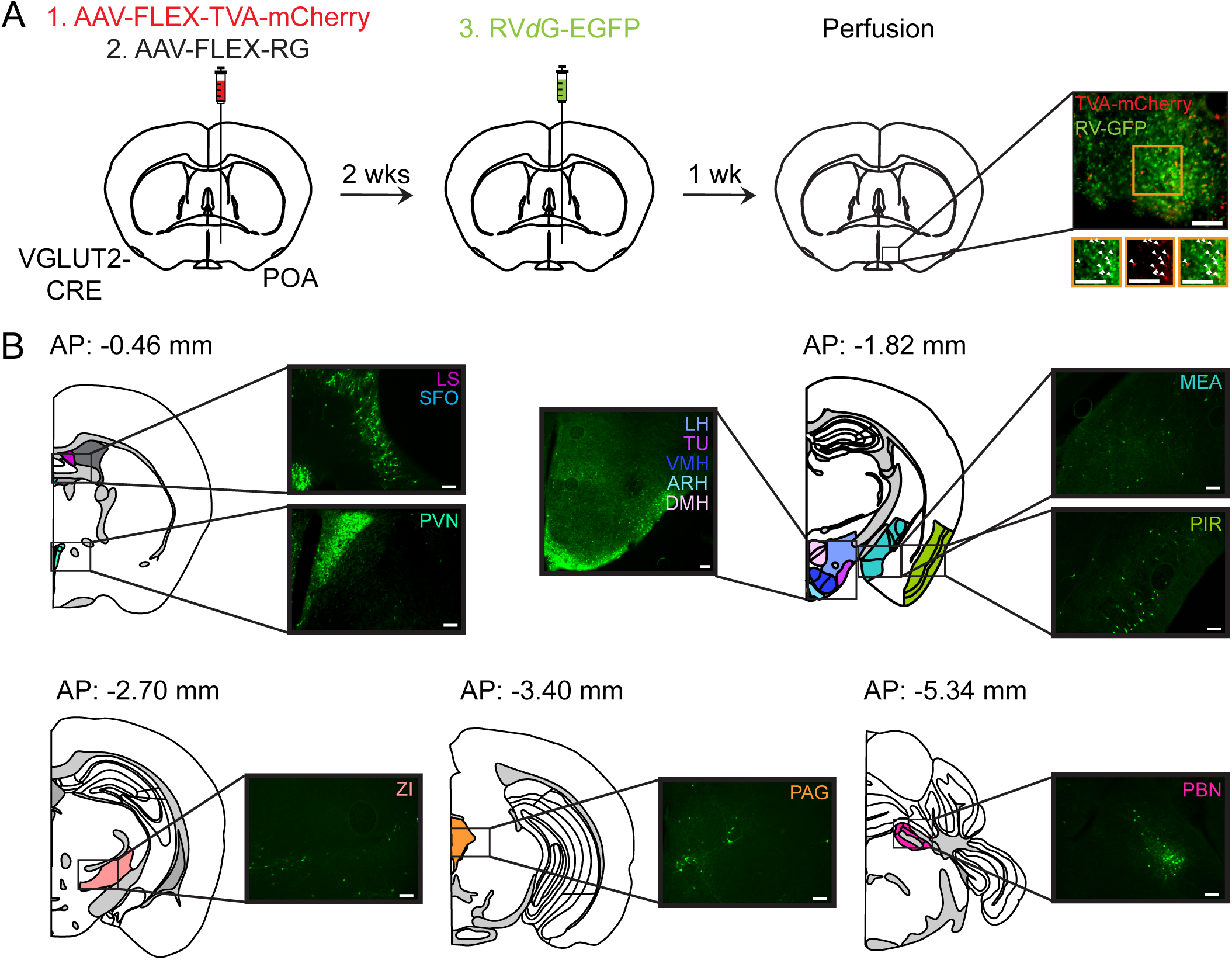
Presynaptic inputs of POA VGLUT2 neurons. **(A)** Schematic illustration of rabies-mediated tracing of monosynaptic inputs to POA VGLUT2 neurons. AAVs expressing a Cre-dependent TVA (the receptor for viruses containing the avian EnvA envelope glycoprotein) fused with mCherry and a Cre-dependent rabies glycoprotein (GP) were injected into the POA of VGLUT2-cre mice. Two weeks later, we injected EnvA-pseudotyped, G-deleted, and GFP expressing rabies virus (RV*d*G-GFP). Starter cells co-express TVA-mCherry and GFP (right). Co-expression of TC and RV*d*G enables rabies viruses to spread to their presynaptic partners. Scale bar, 100 μm **(B)** RV-GFP labeled cells in the lateral septum (LS), subfornical organ (SFO), paraventricular hypothalamic nucleus (PVN), lateral hypothalamus (LH), tuberal nucleus (TU), ventromedial nucleus of the hypothalamus (VMH), arcuate hypothalamic nucleus (ARH), dorsomedial nucleus of the hypothalamus (DMH), medial amygdalar nucleus (MEA), piriform area (PIR), zona incerta (ZI), periaqueductal gray (PAG), and the parabrachial nucleus (PBN). Scale bar, 100 μm

## DISCUSSION

Combining optogenetic manipulation and in vivo calcium imaging with an acute social defeat stress paradigm in mice, we have identified a crucial role of POA glutamatergic neurons in MA regulation during spontaneous sleep and after stress.

### Neural circuits underlying infraslow rhythm and MA regulation

Recently, the activity of LC noradrenergic (LC-NE), dorsal raphe (DR) serotonergic neurons and GABAergic neurons in the dorsomedial medulla has been shown to fluctuate during NREMs in synchrony with the infraslow σ rhythm (Osorio-Forero et al., 2021; Stucynski et al., 2021; Antila et al., 2022; Kato et al., 2022; Kjaerby et al., 2022). The activity dynamics of POA VGLUT2 neurons is strikingly similar to that of LC-NE and DR serotonergic neurons; both populations are negatively correlated with the infraslow σ rhythm and become activated during MAs (Osorio-Forero et al., 2021; Antila et al., 2022; Kato et al., 2022; Kjaerby et al., 2022). Both LC-NE and DR serotonergic neurons innervate the POA (Chou et al., 2002). We and others have previously shown that LC-NE axonal projections to the POA excite neurons that are most active during Wake (Wake-max) and neurons that are active during REMs and wakefulness (Liang et al., 2021; Antila et al., 2022). POA VGLUT2 neurons are most active during wakefulness (**Figure 1**) and, as a subgroup of Wake-max neurons, they are likely excited by LC-NE projections to the POA. Inhibition of LC-NE neurons or their projections to the POA has been shown to decrease the stress-induced increase in MAs (Antila et al., 2022) suggesting that POA VGLUT2 neurons could be a downstream target of LC-NE neurons to regulate MAs and may express α1 or ß adrenergic receptors to mediate the excitatory response of NE. Prazosin, the inverse agonist for α1 receptors, is known to improve sleep and reduce nightmares in PTSD patients (Taylor and Raskind, 2002).

### Role of POA in regulating MAs

During NREMs, POA glutamatergic neurons became activated during MAs, whereas the activity of GABAergic, CCK, and TAC1 neurons was not modulated during MAs (**Figure 1**). Consistent with our findings, a previous study found that chemogenetic activation of POA VGLUT2 neurons promotes sleep fragmentation (Mondino et al., 2021). Optogenetic stimulation of POA VGLUT2 neurons powerfully promotes arousal regardless of the infraslow σ rhythm (**Figure 2**). During undisturbed sleep, animals are less likely to be awakened by external stimuli or to spontaneously wake up during the rising phase of the infraslow σ rhythm, suggesting that this phase constitutes a stable sub-state of NREM sleep. However, while external stimuli are less likely to disturb sleep during this phase, we found that activation of POA VGLUT2 neurons immediately induces arousal, such that the resulting MA or wake episodes become decoupled from the infraslow σ rhythm. Consequently, more frequent activation of POA VGLUT2 neurons as a result of stress will disrupt the rising phase of the infraslow σ rhythm, with adverse effects on sleep continuity. Inhibiting POA VGLUT2 neurons decreased the number of MAs during spontaneous sleep and after stress suggesting that the activity of POA VGLUT2 neurons is necessary for MAs (**Figure 4**). Therefore, the rhythmic fluctuations of POA VGLUT2 neurons may regulate MAs during spontaneous sleep and after stress. In particular, we further reveal that POA VGLUT2 neurons receive monosynaptic inputs from various stress-regulatory regions that could potentially regulate the activity of POA VGLUT2 neurons and consequently sleep quality upon stress exposure (**Figure 5**). A previous study showed that ablation of VGLUT2 neurons in the median preoptic nucleus exacerbated stress-induced insomnia (Machado et al., 2022) suggesting that glutamatergic neurons that are spatially segregated within the preoptic area are differentially involved in stress-induced sleep disturbances. Deleting the NMDA receptor GluN1 subunit from the GABAergic POA neurons was shown to fragment NREMs (Miracca et al., 2022). It remains to be investigated whether POA GABAergic neurons send local inhibitory inputs to the glutamatergic neurons to suppress MAs and consolidate sleep, potentially explaining why deletion of GluN1 subunit in the GABAergic neurons leads to fragmented sleep. Uncovering the brain regions and neural types beyond LC-NE and POA glutamatergic neurons that regulate MAs will help to understand the neural circuit mechanisms underlying sleep fragmentation and, more generally, whether separate circuits are involved in regulating MAs and wakefulness.

Together, our results demonstrate that POA VGLUT2 neurons are an important node in the circuits regulating MAs and thereby directly influence the overall sleep quality after stress. Understanding the circuit mechanisms underlying the regulation of MAs will provide novel therapeutic targets to enhance sleep quality and to alleviate associated symptoms in stress-related sleep disorders.

## METHODS

### Animals

VGLUT2, VGAT, CCK and TAC1-Cre mice (Cat # 016963, 028862, 012706, 021877, Jackson lab) and retired male CD1 breeders (minimum 14 weeks old; Charles River Laboratories) were housed on a 12 h light/12 h dark cycle (lights on 07:00 and off 19:00) with free access to food and water. Male mice were used for social defeat experiments. All the other experiments were performed in males or females. All procedures were approved by Institutional Animal Care and Use Committees of the University of Pennsylvania and were done in accordance with the federal regulations and guidelines on animal experimentation (National Institutes of Health Offices of Laboratory Animal Welfare Policy).

### Surgery

To implant electroencephalogram (EEG) and electromyography (EMG) recording electrodes, adult mice (6 - 12 weeks old) were anesthetized with 1.5 - 2 % isoflurane and placed on a stereotaxic frame. Two stainless steel screws were inserted into the skull 1.7 mm from midline and 1.5 mm anterior to the bregma, and 2 mm from midline and 2 mm posterior to the bregma. The reference screw was inserted on top of the cerebellum. Two EMG electrodes were inserted into the neck musculature. Insulated leads from the EEG and EMG electrodes were soldered to a 2 × 3 pin header, which was secured to the skull using dental cement.

For optogenetic activation experiments, a craniotomy was made on top of the POA in the same surgery as for the EEG and EMG implant, and 0.25 - 0.3 µl of AAV_2_-EF1α-DIO-eYFP or AAV_2_-EF1α-DIO-ChR2-eYFP (University of North Carolina vector core) was injected into the POA (AP 0.4 mm, ML 0.6 mm, DV −5.1 - 5.4 mm from the cortical surface) using Nanoject II (Drummond Scientific) via a micropipette followed by an optic fiber (200 µm in diameter) implantation. For optogenetic inhibition experiments, 0.1 - 0.35 µl of AAV_2_-EF1α-DIO-eYFP or AAV_2_-EF1α-DIO-SwiChR++-eYFP (University of North Carolina vector core) was bilaterally injected into the POA followed by bilateral implantation of optic fibers (200 µm in diameter). Dental cement was applied to cover the exposed skull completely and to secure the optic fiber and the EEG/EMG implant. After surgery, mice were allowed to recover for at least 2-3 weeks before experiments.

For photometry recordings, 0.1 - 0.3 µl of AAV_1_-Syn-Flex-GCaMP6S-WPRE-SV40 (University of Pennsylvania vector core) was injected into POA and the optic fiber (400 µm in diameter) was placed on top of the injection site.

For rabies tracing experiments, 0.3 - 0.4 µl of a mixture containing AAV-CAG-FLEX-TVA–mCherry and AAV-CAG-FLEX-RG was injected into the POA. After 2 weeks, 0.4 - 0.45 µl of EnvA-pseudotyped, rabies-glycoprotein-deleted, and GFP-expressing rabies viral particles (RV*dG*) were injected into the POA. After 1 week, mice were perfused. Rabies tracing viruses were obtained from the University of California, Irvine.

### Histology

Mice were deeply anesthetized and transcardially perfused with phosphate-buffered saline (PBS) followed by 4 % paraformaldehyde (PFA) in PBS. Brains were fixed overnight in 4% PFA and then transferred to 30 % sucrose in PBS solution for at least one night. Brains were embedded and mounted with Tissue-Tek OCT compound (Sakura Finetek) and frozen. 40-60 μm sections were cut using a cryostat (Thermo Scientific HM525 NX) and mounted onto glass slides. Brain sections were washed with PBS followed by counterstaining with Hoechst solution (#33342, Thermo Scientific). Slides were cover-slipped with Fluoromount-G (Southern Biotechnic) and imaged using a fluorescence microscope (Microscope, Leica DM6B; Camera, Leica DFC7000GT; LED, Leica CTR6 LED).

### Sleep recordings

Sleep recordings were carried out in the animal’s home cage or in a cage to which the mouse had been habituated. EEG and EMG electrodes were connected to flexible recording cables via a mini-connector. EEG and EMG signals were recorded using an RHD2132 amplifier (Intan Technologies, sampling rate 1 kHz) connected to the RHD USB Interface Board (Intan Technologies). For fiber photometry, we used a Tucker-Davis Technologies RZ5P amplifier (sampling rate 1.5 kHz). EEG and EMG signals were referenced to a ground screw placed on top of the cerebellum. To determine the brain state of the animal, we first computed the EEG and EMG spectrogram for sliding, half-overlapping 5 s windows, resulting in 2.5 s time resolution. To estimate within each 5 s window the power spectral density (PSD), we performed Welch’s method with Hanning window using sliding, half-overlapping 2 s intervals. Next, we computed the time-dependent δ (0.5 to 4 Hz), θ (5 to 12 Hz), σ (12 to 20 Hz) and high γ (100 to 150 Hz) power by integrating the EEG power in the corresponding ranges within the EEG spectrogram. We also calculated the ratio of the θ and δ power (θ/δ) and the EMG power in the range 50 to 500 Hz. For each power band, we used its temporal mean to separate it into a low and high part (except for the EMG and θ/δ ratio, where we used the mean plus one standard deviation as threshold). REMs was defined by a high θ/δ ratio, low EMG, and low δ power. A state was set as NREMs if the δ power was high, the θ/δ ratio was low, and EMG power was low. In addition, states with low EMG power, low δ, but high σ power were scored as NREMs. Wake encompassed states with low δ power and high EMG power and each state with high γ power (if not otherwise classified as REMs). Our automatic algorithm that has been published in (Weber et al., 2015, 2018; Chung et al., 2017; Stucynski et al., 2021; Antila et al., 2022) has 90.256 % accuracy compared with the manual scoring by expert annotators. We manually verified the automatic classification using a graphical user interface visualizing the raw EEG and EMG signals, EEG spectrograms, EMG amplitudes, and the hypnogram to correct for errors, by visiting each single 2.5 sec epoch in the hypnograms. The software for automatic brain state classification and manual scoring was programmed in Python (available at https://github.com/tortugar/Lab/tree/master/PySleep). For **Figure 1D**, we visually detected the onset of MAs using 0.25 sec precision as events characterized by a desynchronized EEG accompanied by motor-bursts.

### Optogenetic manipulation

Light pulses (for activation: 4 s pulse, 10 Hz, at 15 - 25 min interval, 2 - 3.5 mW; for inhibition: 3 s step pulses at 60 s intervals, 2 - 4 mW) were generated by a blue laser (473 nm, Laserglow) and sent through the optic fiber (200 µm diameter, ThorLabs) that connects to the ferrule on the mouse head. TTL pulses to trigger the laser were controlled using a raspberry pi, which was controlled by a custom user interface programmed in Python. Optogenetic manipulations were conducted during the light period for 8 - 10 hrs (activation) or 4 hrs (inhibition).

### Fiber photometry

For calcium imaging, a first LED (Doric lenses) generated the excitation wavelength of 465 nm and a second LED emitted 405 nm light, which served as control for bleaching and motion artifacts. The 465 and 405 nm signals were modulated at two different frequencies (210 and 330 Hz). Both lights were passed through dichroic mirrors before entering a patch cable attached to the optic fiber. Fluorescence signals emitted by GCaMP6s were collected by the optic fiber and passed via the patch cable through a dichroic mirror and GFP emission filter (Doric lenses) before entering a photoreceiver (Newport Co.). Photoreceiver signals were relayed to an RZ5P amplifier (Tucker-Davis Technologies, TDT) and demodulated into two signals using TDT’s Synapse software, corresponding to the 465 and 405 nm excitation wavelengths. To analyze the calcium activity, we used custom-written Python scripts. First, both signals were low-pass filtered at 2 Hz using a 4th order digital Butterworth filter. Next, using linear regression, we fitted the 405 nm to the 465 nm signal. Finally, the linear fit was subtracted from the 465 nm signal (to correct for photo-bleaching or motion artifacts) and the difference was divided by the linear fit yielding the ΔF/F signal. To determine the brain state, EEG and EMG signals were simultaneously recorded with calcium signals using the RZ5P amplifier.

To detect calcium transients occuring on the infraslow timescale, we first filtered the calcium signal with a zero-lag, 4th order digital Butterworth filter with cutoff frequency 1/15 Hz. Next, we detected prominent peaks in the signal using the function find_peaks provided by the python library scipy (https://scipy.org/). As parameter for the peak prominence, we used 0.05 * distance between the 1st and 99th percentile of the distribution of the ΔF/F signal.

### Acute social defeat stress

We followed published social defeat stress procedures (Golden et al., 2011; Antila et al., 2022) with slight modifications. Retired male CD1 breeders were initially screened for aggressiveness on 3 consecutive days. Mice initiating an aggressive attack toward a previously unknown intruder mouse in 10 - 60 s on at least 2 consecutive days were used as aggressors. In the acute social defeat paradigm, a CD1 mouse was housed on one side of a cage, which was partitioned with a plastic perforated divider. The experimental mouse was placed directly within the CD1’s home cage compartment for 5 - 10 min until 10 bouts of physical aggression by the CD1 were reached. Subsequently, the aggressor mouse was removed and the experimental mouse was transferred to the opposite side of the divider during which the sleep recordings were conducted. Social defeat was always performed at ZT 3 - 5. Each mouse was exposed to the defeat once. To limit potential physical pain, mice were exposed to the defeat for less than 10 minutes and maximally 10 attacks (whichever comes first), and specific aggressors from the experiment were removed if they were causing consistent wounding similar to a widely used protocol (Golden et al., 2011). Furthermore, we carefully observed the mice during the defeat, and prolonged attacks by CD1 mice were interrupted to prevent physical injuries to the experimental mouse. In case a mouse suffered from pain, it was excluded from our study.

### Analysis of infraslow σ power oscillations

To calculate the power spectral density of the EEG σ power, we first calculated for each recording the EEG power spectrogram by computing the FFT for consecutive sliding, half-overlapping 5 s windows. Next, we normalized the spectrogram by dividing each frequency component by its mean power and calculated the normalized σ power by averaging across the spectral density values in the σ range (10.5 - 16 Hz). As the infraslow rhythm is most pronounced in consolidated NREMs bouts (Lecci et al., 2017), we only considered NREMs bouts that lasted at least 120 s, possibly interrupted by MAs (wake periods ≦ 20s). We then calculated the power spectral density using Welch’s method with Hanning window for each consolidated NREMs bout and averaged for each animal across the resulting densities. To quantify the strength of the infraslow rhythm, we computed the areas under the PSD in the ranges 0.01 - 0.04 Hz and 0.08 - 0.12 Hz, respectively, and subtracted the second value from the first value.

To determine the instantaneous phase of the infraslow rhythm, we smoothed the σ power using a 10 s box filter and band-pass filtered it in the range of 0.01 - 0.03 Hz using a 4th order digital Butterworth filter. Finally, we computed the phase angle by applying the Hilbert transform to the band-pass filtered σ power signal. Based on the phase, we could then isolate the beginning and end of single infraslow cycles to average the ΔF/F activity during single cycles.

To calculate the cross-correlation between the ΔF/F signal and the σ power (or other power bands), we first calculated for all NREMs bouts with duration ≧ 120 s (possibly interrupted by MAs) the σ power (s) from the EEG spectrogram, computed using consecutive 2.5 s windows with 80 % overlap to increase the temporal resolution. We again normalized the spectrogram by dividing each frequency component by its mean power. Using the same overlapping binning, we downsampled the ΔF/F signal (d) to prevent any time lags resulting from differences in downsampling and then calculated the cross-correlation of both signals. The cross-correlation was normalized by dividing it by the product of the standard deviation of s and d and the number of data points in s. For each mouse we finally obtained the mean cross-correlation by averaging across all NREMs bouts.

### Sleep spindle detection

Spindles were detected using a previously described algorithm using the frontal EEG (Park et al., 2021). The spectrogram was computed for consecutive 600 ms windows with 500 ms overlap, resulting in a 100 ms temporal resolution. The spindle detection algorithm used two criteria to determine for each 100 ms time bin whether it was part of a spindle or not: The first criterion was that the height of the maximum peak in the σ frequency range (10 - 16.67 Hz) exceeds a threshold, which corresponded to the 96th percentile of all maximum peaks in the σ frequency range of the sleep recording. We determined the optimal percentile value by maximizing the performance of the algorithm on a manually annotated control data set. Second, the power value of the peak in the σ range (10 - 16.67 Hz) had to be greater than half of the peak value in the range 0 - 10 Hz. The optimal value for this ratio (σ peak ratio) was again determined on the control data set. Next, the algorithm merged spindle events that were temporally close to each other. First, spindle events in adjacent bins were considered as part of the same spindle. Second, we fused together sequences of spindle events that were interrupted by gaps of less than 300 ms. The optimal value for the gap was again determined on the control data set. Finally, we discarded spindles with duration ≦ 200 ms. Of all the potential spindles, we only considered those as spindles where for at least half of the time bins the peak frequency lied in the range of 10 - 16.7 Hz. The parameters of the spindle detection algorithm (σ percentile threshold, σ peak ratio, and minimum fusing distance) were optimized using a manually annotated data set.

### Statistical tests

Statistical analyses were performed using the python packages scipy.stats (scipy.org) and pingouin (https://pingouin-stats.org) (Vallat, 2018). We did not predetermine sample sizes, but cohorts were similarly sized as in other relevant sleep studies. All statistical tests were two-sided. Data were compared using t-tests or ANOVA followed by multiple comparisons tests. To account for multiple comparisons, p-values were Bonferroni corrected. For all tests, a (corrected) p-value < 0.05 was considered significant. Box plots were used to illustrate the distribution of data points. The upper and lower edges of the box correspond to the quartiles (25th and 75th percentile) of the dataset and the horizontal line in the box depicts the median, while the whiskers indicate the remaining distribution, except for outliers, i.e. points smaller than the 25th percentile −1.5 * the interquartile range (IQR) or larger than the 75th percentile + 1.5 IQR. Outliers are depicted as diamonds. In **Figure 1D**, we first used one-way rm ANOVA to determine whether the activity was significantly modulated throughout this time interval. If it is significant, we used consecutive pairwise t tests with Bonferroni correction to determine when the activity starts to become significantly modulated. Statistical results are presented in the text.

## ACKNOWLEDGMENTS

This work was supported by the National Institute of Neurological Disorders and Stroke (R01-NS-110865), the Whitehall Foundation, Alfred P. Sloan Foundation, a NARSAD Young Investigator grant, Simons Foundation Pilot Award, Eagle Autism Challenge Pilot Grant, The McCabe Fund Award, The Hartwell Individual Biomedical Research Award to S. C. We thank the members from Chung and Weber labs for helpful discussion.

## AUTHOR CONTRIBUTIONS

Conceptualization, J.S., and S.C.; Methodology, J.S., H.A. and K.B.; Software, H.A., F.W.; Investigation, J.S., A.H.F.; Writing - Original Draft, J.S. and S.C.; Writing - Review & Editing, J.S., F.W., S.C.; Supervision, S.C.; Funding Acquisition, S.C.

## DECLARATION OF INTERESTS

The authors declare no competing financial interests.

---

**Figure S1.**
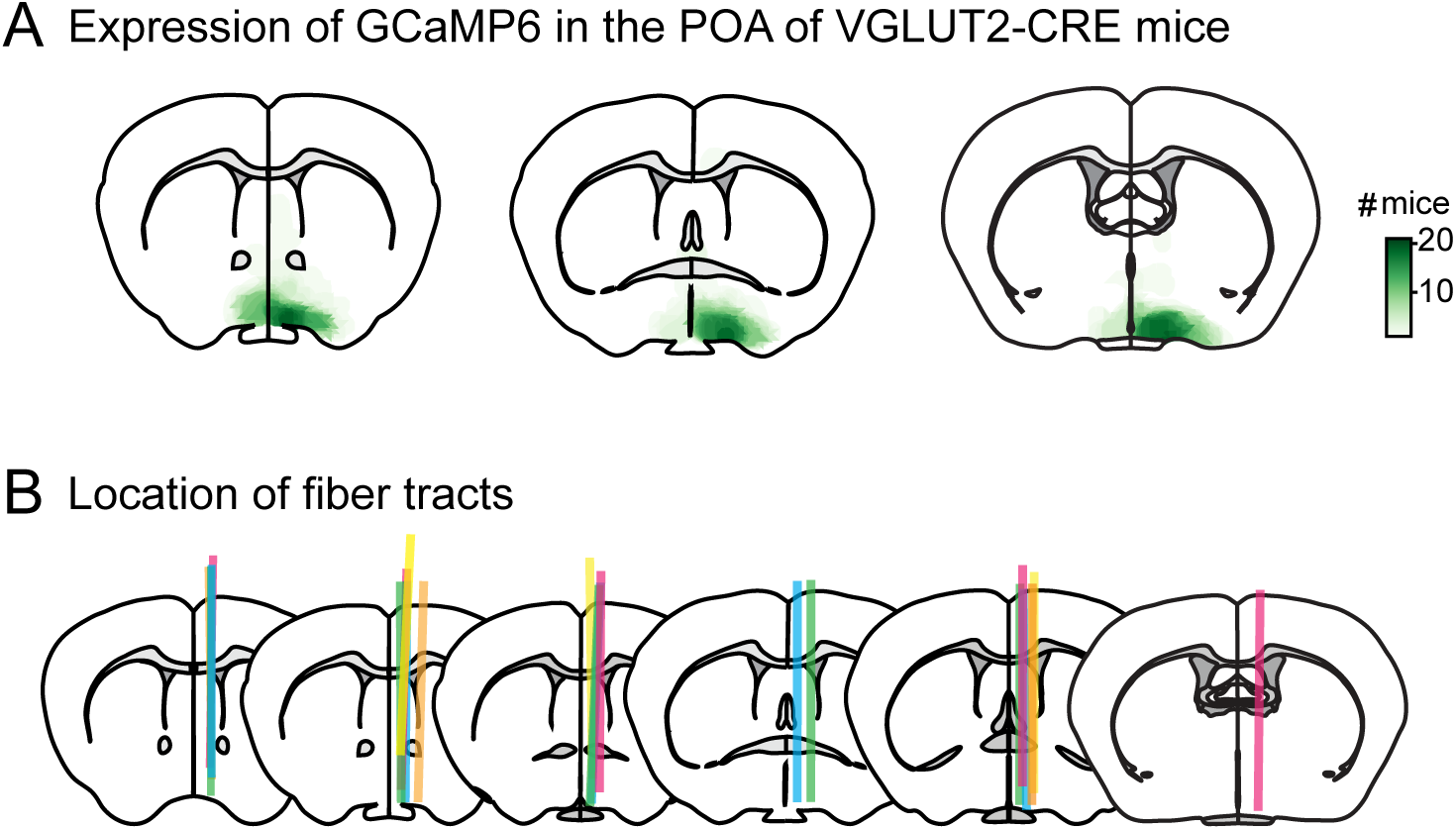
Heatmaps of GCaMP6 expression and location of optic fibers in the POA of VGLUT2-cre photometry mice, related to Figure 1. **(A)** Heatmaps depicting expression of GCaMP6s in the POA of VGLUT2-cre mice. n = 21 mice. **(B)** Location of fiber tracts. Each colored bar represents the location of optic fibers for photometry recordings.

**Figure S2.**
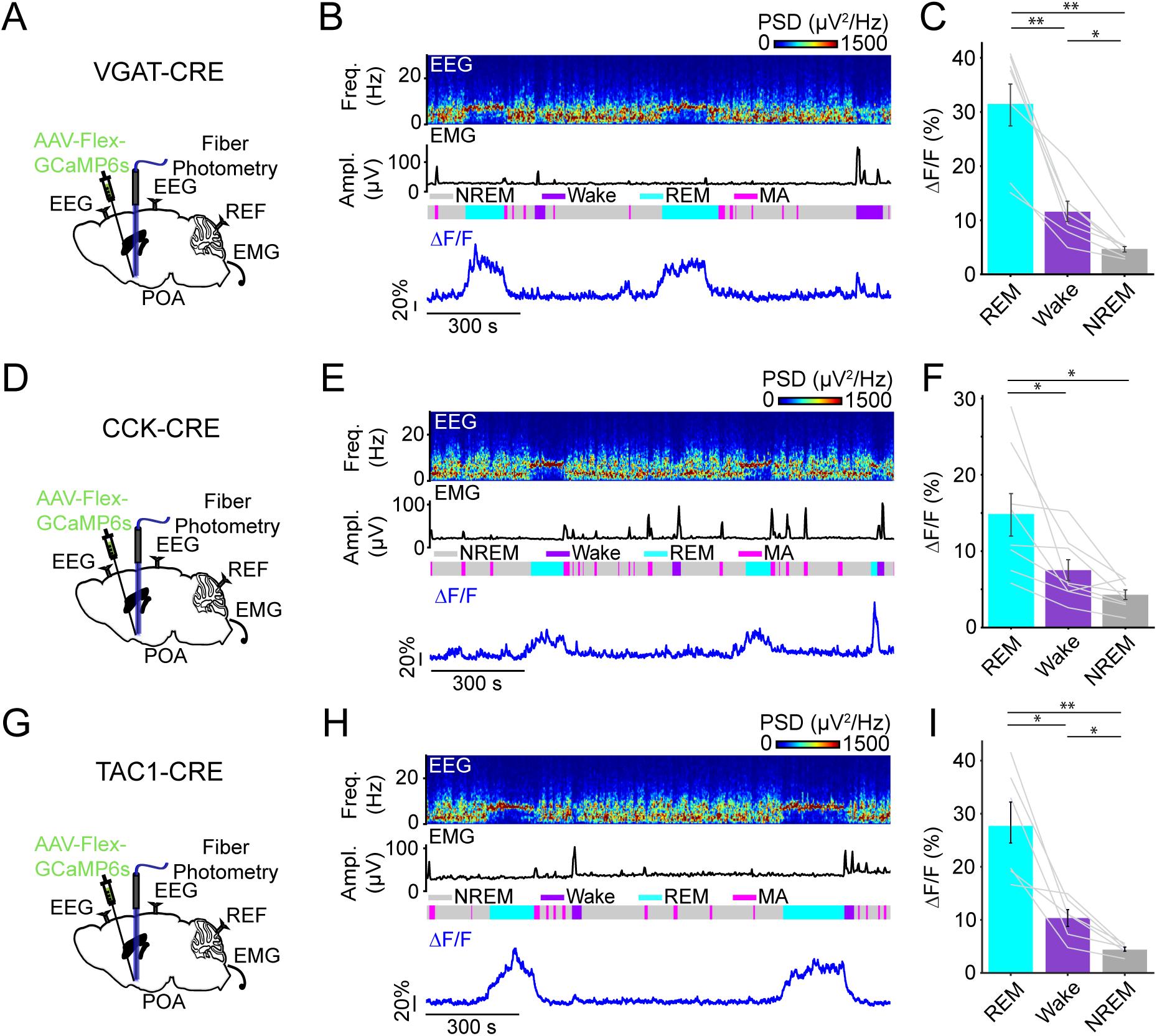
POA VGAT, CCK and TAC1 neurons are most active during REMs, related to Figure 1. **(A)** Schematic of fiber photometry in POA VAGT neurons with simultaneous EEG and EMG recordings. **(B)** Example fiber photometry recording of POA VGAT neurons. Shown are parietal EEG spectrogram, EMG amplitude, color-coded brain states, and ΔF/F signal. **(C)** ΔF/F activity during REMs, wake, and NREMs. n = 7 mice. **(D)** Schematic of fiber photometry in POA CCK neurons with simultaneous EEG and EMG recordings. **(E)** Example fiber photometry recording of POA CCK neurons. **(F)** ΔF/F activity during REMs, wake, and NREMs. n = 8 mice. **(G)** Schematic of fiber photometry in POA TAC1 neurons with simultaneous EEG and EMG recordings. **(H)** Example fiber photometry recording of POA TAC1 neurons. **(I)** ΔF/F activity during REMs, wake, and NREMs. n = 6 mice. **(C, F, I)** Bars, averages across mice; lines, individual mice; error bars, ± s.e.m. One-way rm ANOVA followed by pairwise t tests with Bonferroni correction, **P < 0.01; *P < 0.05.

**Figure S3.**
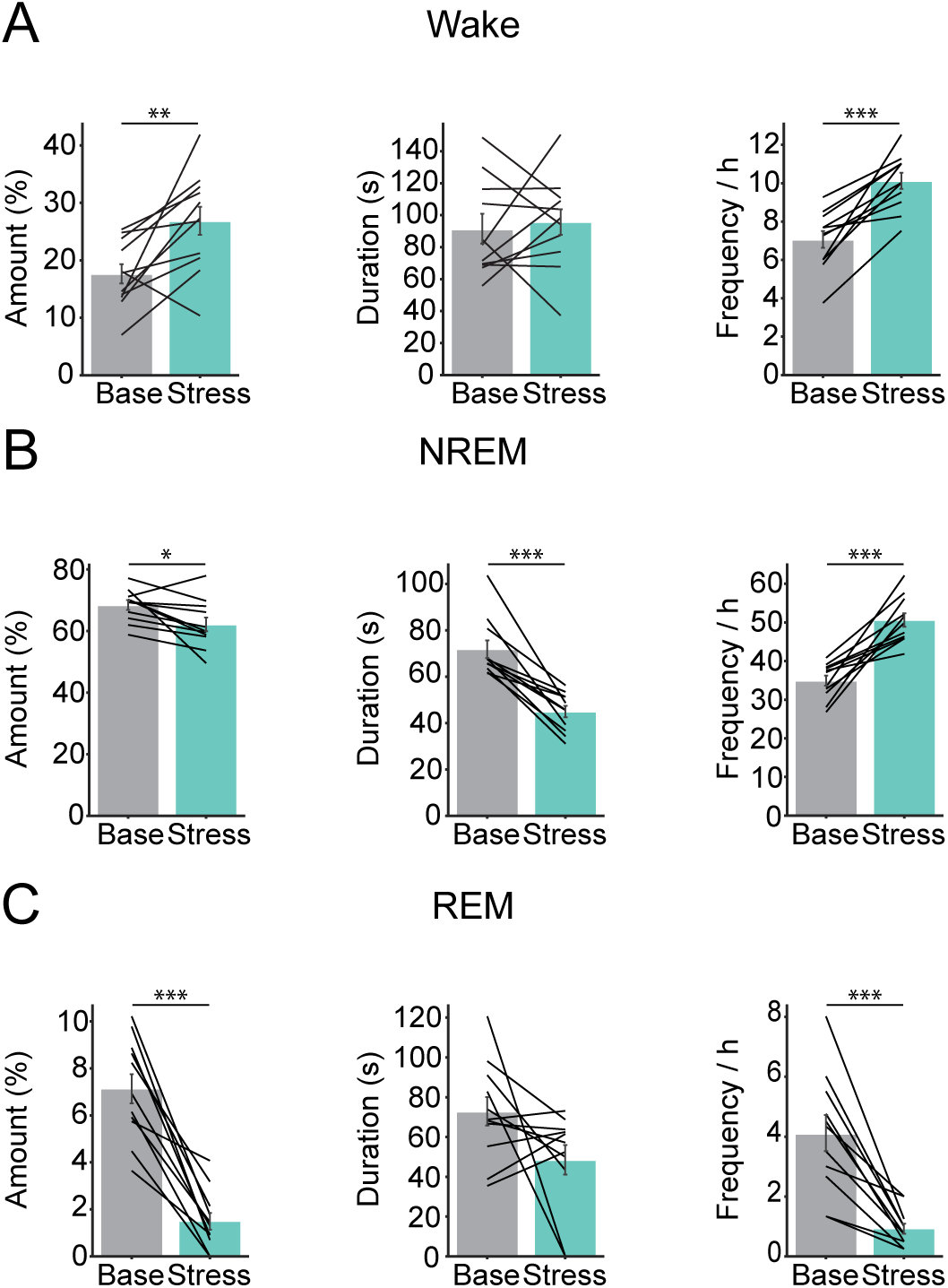
Impact of acute social defeat stress on brain states. Related to Figure 3. **(A)** Percentage of time in wake, duration and frequency of wake episodes. **(B)** Percentage of time in NREMs, duration and frequency of NREMs episodes. **(C)** Percentage of time in REMs, duration and frequency of REMs episodes. Bars, averages across mice; lines, individual mice; error bars, ± s.e.m. Paired t tests, ***P < 0.001; **P < 0.01; *P < 0.05. n = 11 mice.

**Figure S4.**
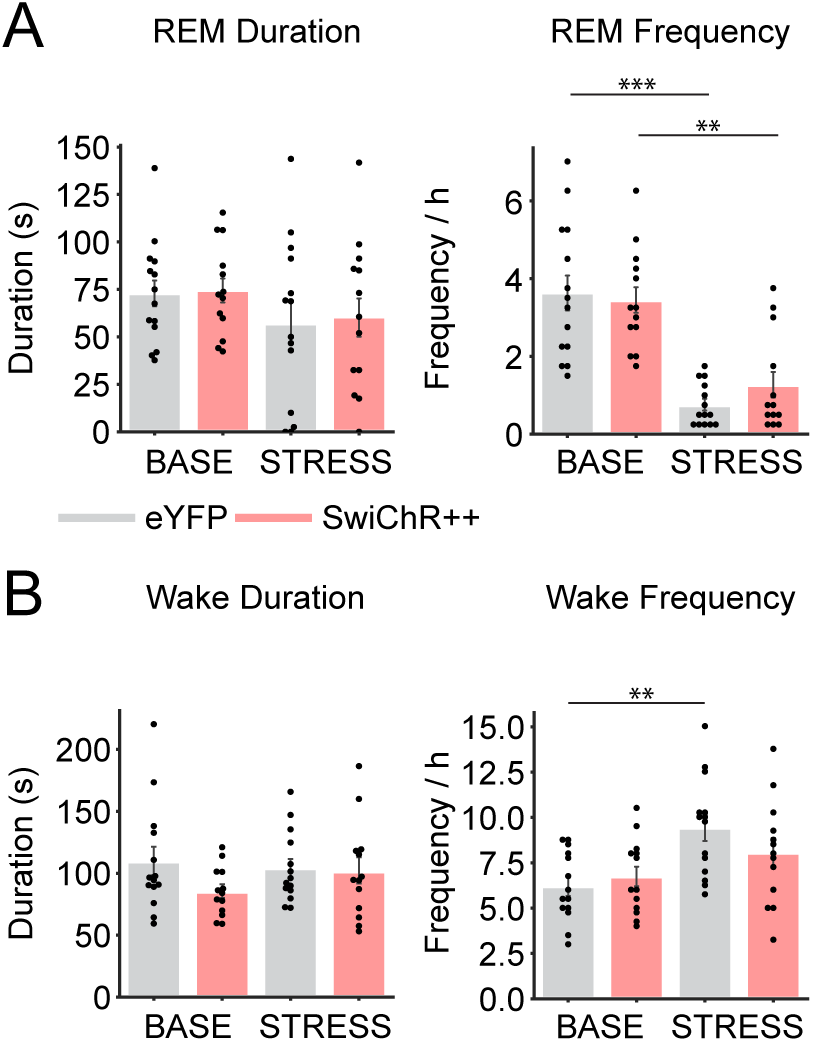
Effect of SwiChR++-mediated inhibition of POA VGLUT2 neurons on duration and frequency of REMs and wake episodes, related to Figure 4. **(A)** Duration and frequency of REMs episodes during laser baseline and stress recordings. **(B)** Duration and frequency of wake episodes during laser baseline and stress recordings. Bars, averages across mice; dots, individual mice; error bars, ± s.e.m. Mixed ANOVA followed by pairwise t-tests with Bonferroni correction, ***P < 0.001; **P < 0.01. n = 13-14 mice.

## Notes

### Competing Interest Statement

The authors have declared no competing interest.

### Summary of Updates

Added new figure related to optogenetic activation and updated related text; added new figure related to tracing experiments and updated related text; revised figures; revised text; added supplemental figures

